# Understanding the compositional changes in LB Lennox medium during growth of *Escherichia coli* DH5α in a 1L bioreactor

**DOI:** 10.1101/2020.09.24.311373

**Authors:** Wenfa Ng

**Affiliations:** Department of Chemical and Biomolecular Engineering, National University of Singapore

**Keywords:** gel filtration chromatography, reversed phase high performance liquid chromatography, bioreactor, growth medium, fractions, chromatogram, wavelength of visualization

## Abstract

Compositional changes in growth medium represents dynamic interplay between cell growth, biomass formation, and energy maintenance, with concomitant decrease in nutrients and increase in secreted metabolites and metabolic byproducts. Such information is important for quantifying microbial physiological response both at the population and cellular level, with respect to understanding subtle differences in microbial growth response, as well as supporting model building efforts in metabolic engineering. With the desire to understand molecular weight changes in components of LB Lennox medium as well as broth fractions present during cultivation of *Escherichia coli* DH5α (ATCC 53868) at 37 °C, 400 rpm stirring and 1 VVM aeration in a 1 L bioreactor, this study used a combination of gel filtration chromatography (GFC) and reversed phase high performance liquid chromatography (RP-HPLC) for determining changes to molecular weight of different fractions of the growth medium. Experiment results revealed the difficulty of fractionating the culture broth with RP-HPLC, where no distinct peaks of narrow retention time width were obtained. More importantly, the column used for GFC was unable to differentiate small molecular weight changes on the order of a few tens to few hundred Da through a refractive index detector. Together, GFC and RP-HPLC highlighted the difficulty of fractionating LB Lennox culture broth into different distinct fractions. Finally, the study validated the use of 194 nm as detection wavelength for visualizing the chromatogram of LB Lennox medium eluted from a C-18 reversed phase column during liquid chromatography. Collectively, GFC and RP-HPLC could not fractionate LB Lennox broth of *E. coli* DH5α into distinct fractions for further analysis by identification techniques such as mass spectrometry. Given the inherent complexity of complex medium such as LB Lennox, clean separation of the medium into every component with high purity may be impossible.

**Graphical abstract:** 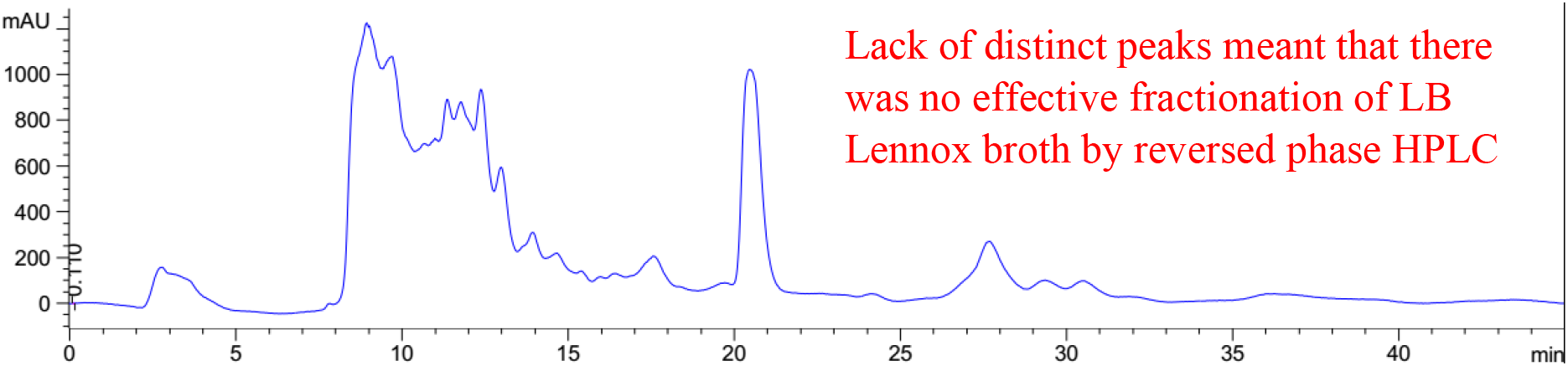

**Short description:** Broad elution profile and lack of distinct peaks in chromatogram of LB Lennox broth at 6 hours post inoculation with *Escherichia coli* DH5α (ATCC 53868), after attempted fractionation by C-18 reversed phase high performance liquid chromatography (RP-HPLC), revealed that separation could not be achieved with a complex mixture such as a microbiology broth. Specifically, while the objective of understanding compositional changes to culture medium during growth presents tremendous opportunities for determining the dynamic conversion of nutrients into metabolites and byproducts, difficulty of profiling all chemical constituents in a complex broth mixture meant that current model building efforts for metabolic engineering remains primitive with respect to the consortia of metabolic reactions occurring *in situ* at the cellular level.

**Subject areas:** biotechnology, biochemistry, cell biology, microbiology, analytical chemistry

## 1. Introduction

Composition of growth medium changes dynamically as a microorganism grows, and many facets of metabolism and metabolic flow in a cell remains enigmatic due to the inability to understand the changes in growth medium composition during cultivation, especially in the case of complex undefined medium. Such information is important to understanding how different nutrients contribute to energy maintenance and biomass formation in a cell,^1,2^ where energy derived from various nutrients are partitioned into different metabolic pathways at the cellular level, that in aggregate, manifest as detectable changes in the medium composition at the cell population level.

Determination of small molecule metabolites such as glucose, glycerol, acetate, fructose, xylose, and lactate is easier, and could readily be achieved through a chromatography column with combined reversed phase and ion exchange separation capability.^3^ An important unsolved problem in cellular metabolism remains the determination of compositional changes to growth medium, especially complex medium, where significant difficulty exists in understanding, in totality, all components present. Specifically, the multi-dimensional space of nutrient composition in a growth medium, and its dynamic changes with depletion of nutrients and infusing of metabolic products of cells meant that characterizing the compositional changes by instrumented techniques such as high performance liquid chromatography mass spectrometry (LC-MS) remains difficult. The key problem is the inability to fractionate the mixture into individual components for mass spectrometry detection.^4^ Thus, while modern tandem mass spectrometry techniques have afforded the ability to determine the molecular weight of individual species in a sample or fraction,^5,6^ the challenge lies in developing a separation protocol that would yield a pure sample of individual component for analysis.

On the other hand, multiple dimensions of separation are necessary to fractionate and quantify the diverse mix of carbohydrates, proteins, peptides, amino acids, lipids and other small molecules present in the culture broth. This meant that multiple types of columns with different chemistries must be concatenated in sequence to effect separation of a complex medium broth. Overall, understanding the compositional changes in a growth medium during microbial cultivation remains difficult, and awaits technological breakthroughs in instrumented techniques. But, current chromatography instrumentations may be able to lend some insights into the major fractions in a culture broth, and help provide some initial understanding of the complex multi-dimensional problem.

Seeking to understand the compositional changes to LB Lennox medium as *Escherichia coli* DH5α (ATCC 53868) grows in it, gel filtration chromatography (GFC) and C-18 reversed phase high performance liquid chromatography (RP-HPLC) were used to characterize the changes to the culture broth with growth of the bacterium. The objective was to determine changes to the growth medium at the molecular weight fraction level and composition level. To this end, *E. coli* DH5α was grown in LB Lennox (unbuffered) medium in a 1L bioreactor in a fed-batch mode, where glucose and NH_4_Cl were infused to the bioreactor after the culture had entered stationary phase. Samples collected at regular time points in the culture were analysed by both gel filtration chromatography for understanding the molecular weight changes in the different fractions present, as well as RP-HPLC for determining the different fractions.

Results revealed that the gel filtration column used was unable to detect molecular weight changes of a few Da to few hundred Da. Three broad peaks were seen in the eluted chromatogram from gel filtration chromatography, indicating that the column was unable to distinguish fractions beyond the three broad ones detected. Reversed phase high performance liquid chromatography, on the other hand, was also unable to fractionate LB Lennox broth given the complexity of the microbiological medium. Additionally, 194 nm was validated to be a useful wavelength for visualizing the RP-HPLC chromatogram of LB Lennox medium given the significant increase in information available at this wavelength compared to 280, 264 and 254 nm.

## 2. Materials and methods

### 2.1 Materials

LB Lennox medium was purchased from Difco and has the following composition [g/L]: Tryptone, 10.0; Yeast Extract, 5.0; NaCl, 5.0. Polyethylene glycol antifoam was purchased from Sigma-Aldrich.

### 2.2 Growth of *Escherichia coli* DH5α in bioreactor

Glycerol stock culture of *Escherichia coli* DH5α (ATCC 53868), kept at −70 °C prior to inoculation, was used in inoculating a 100 mL LB Lennox medium in a 250 mL glass shake flask. Cultivation of the aerobic seed culture was at 37 °C and 230 rpm rotational shaking in a temperature controlled incubator. After 14 hours of incubation, 5 mL of the seed culture was used in inoculating 500 mL of LB Lennox medium and 50 mL of 100 g/L polyethylene glycol antifoam. Stirring speed was 400 rpm with 1 VVM aeration. Temperature was controlled at 37 °C, with no control for pH. At appropriate timepoints, 5 mL of broth was collected for analysis of pH and optical density.

Optical density was measured with a Shimadzu Biospec-Mini spectrophotometer using a quartz cuvette of 1 cm pathlength, where appropriate dilution with deionized water was used if optical density exceeded 1. pH was measured with an Orion pH probe fitted on a Sartorius pH meter.

### 2.3 Gel filtration chromatography (GFC)

Waters gel filtration chromatography instrument was calibrated with polyethylene glycol molecular weight standards prior to the experiment. Refractive index detector was used for detection, and the column and detector temperatures were at 35 °C. 0.1g of the molecular weight standards were dissolved in 100 mL of ultrapure water one day prior to the analysis and the constituted standards filtered through 0.45 μm nylon membrane filter. Molecular weight markers used were 1460 Da, 7130 Da, 12900 Da, and 20600 Da.

Samples from the bioreactor were filtered through 0.22 μm nylon membrane filters prior to injection into an Agilent Technologies PL aquagel-OH MIX 8 μm gel filtration column with an injection volume of 10 μL. Flow rate for elution was 1 ml/min with water as eluent.

### 2.4 Reversed phase high performance liquid chromatography (RP-HPLC)

LB Lennox broth samples were filtered through 0.22 μm nylon membrane filters prior to injection into an Agilent Technologies Poroshell 120SB C-18 column attached to an Agilent 1200 series high performance liquid chromatography (HPLC) instrument. Injection volume was 10 μL, and detector used was a diode array detector.

## 3. Results and Discussion

*Escherichia coli* DH5α (ATCC 53868) was grown in LB Lennox medium (unbuffered) medium at 37 °C in a 1 L bioreactor with 1 VVM of aeration and 10 g/L of polyethylene glycol antifoam (Figure 1). Specifically, *E. coli* DH5α shown good growth in the medium with optical density of 3.2 at 6 hours post inoculation. pH decreased from 6.9 at the start of the culture to 6.5 after 2 hours of cultivation, but increased thereafter to 7.8 at 6 hours of cultivation. Changes in the pH profile between 2 and 6 hours post inoculation revealed that metabolic byproducts could have been secreted by the cells into the culture broth.

**Figure 1:**
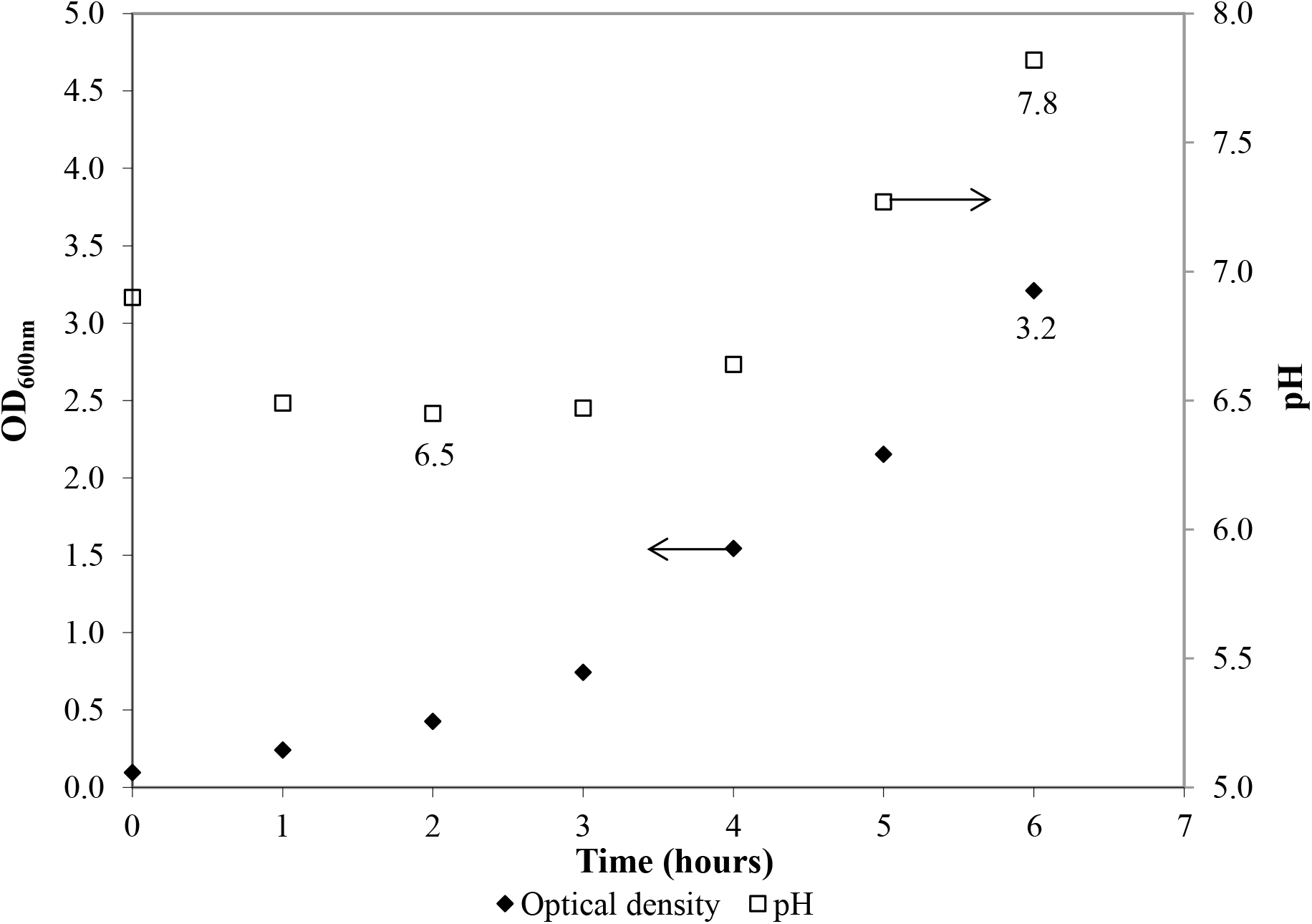
Growth of *Escherichia coli* DH5α in LB Lennox (unbuffered) medium at 37 °C in a 1 L bioreactor with aeration = 1 VVM, and 10 g/L polyethylene glycol antifoam.

Attempts to determine the various fractions that existed in LB Lennox broth during a time course experiment did not succeed in identifying beyond three major fractions of the broth that existed throughout the 6 hours experiment (Figure 2 to 8). Specifically, gel filtration chromatography could not fractionate the different fractions present in LB Lennox broth. More importantly, given that the elution profile of the 3 major peaks did not shift substantially during the experiment, gel filtration chromatography coupled with the Agilent PL aquagel-OH MIX 8 μm column did not separate the complex LB Broth mixture into more than 3 fractions.

**Figure 2:**
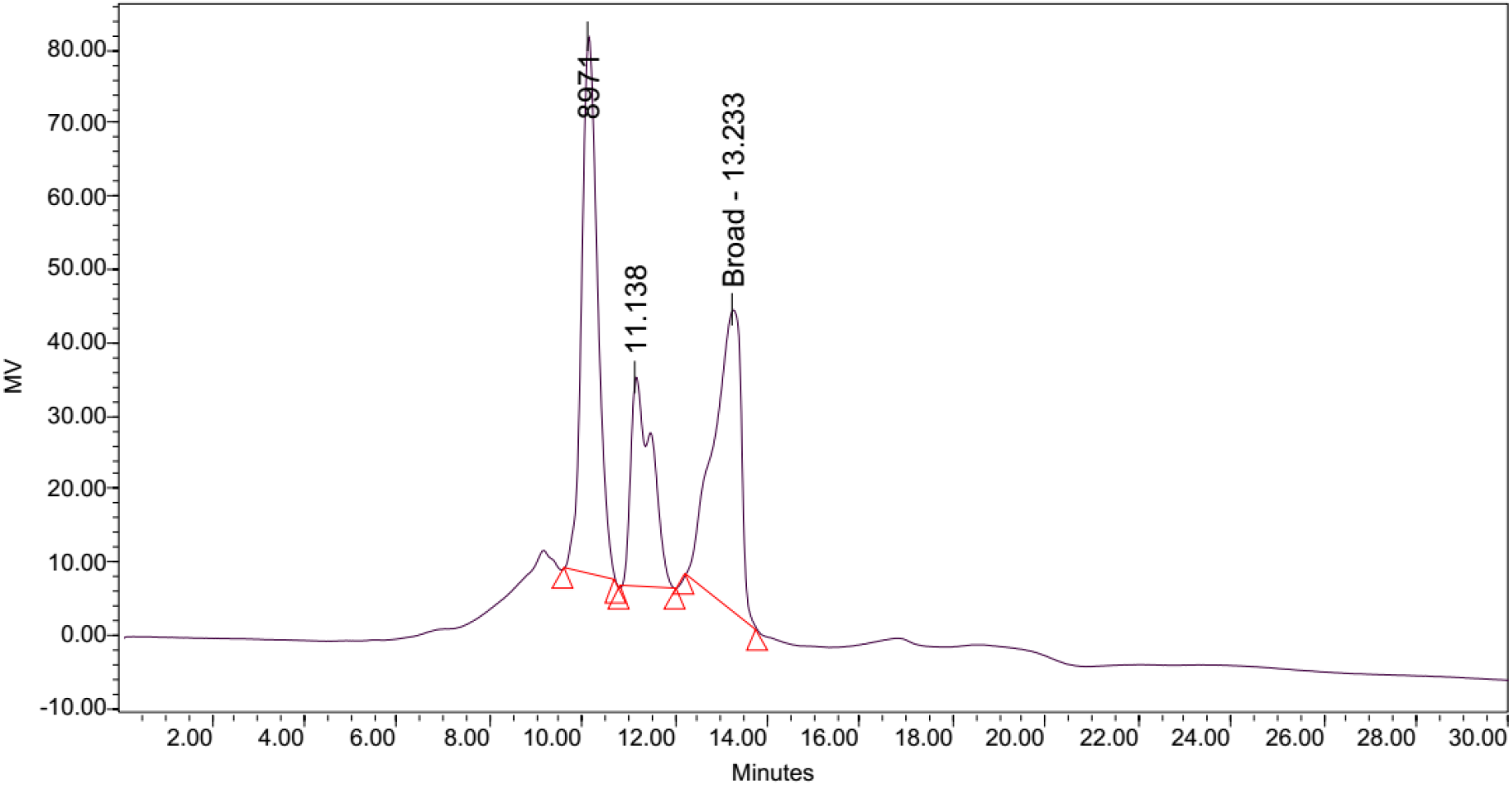
Gel filtration chromatography separation of LB Lennox broth at time = 0 hrs reveals the existence of three peaks. Injection volume = 10 μL, flow rate = 1 ml/min. Column: Agilent PL aquagel-OH MIX 8 μm, refractive index detector

**Figure 3:**
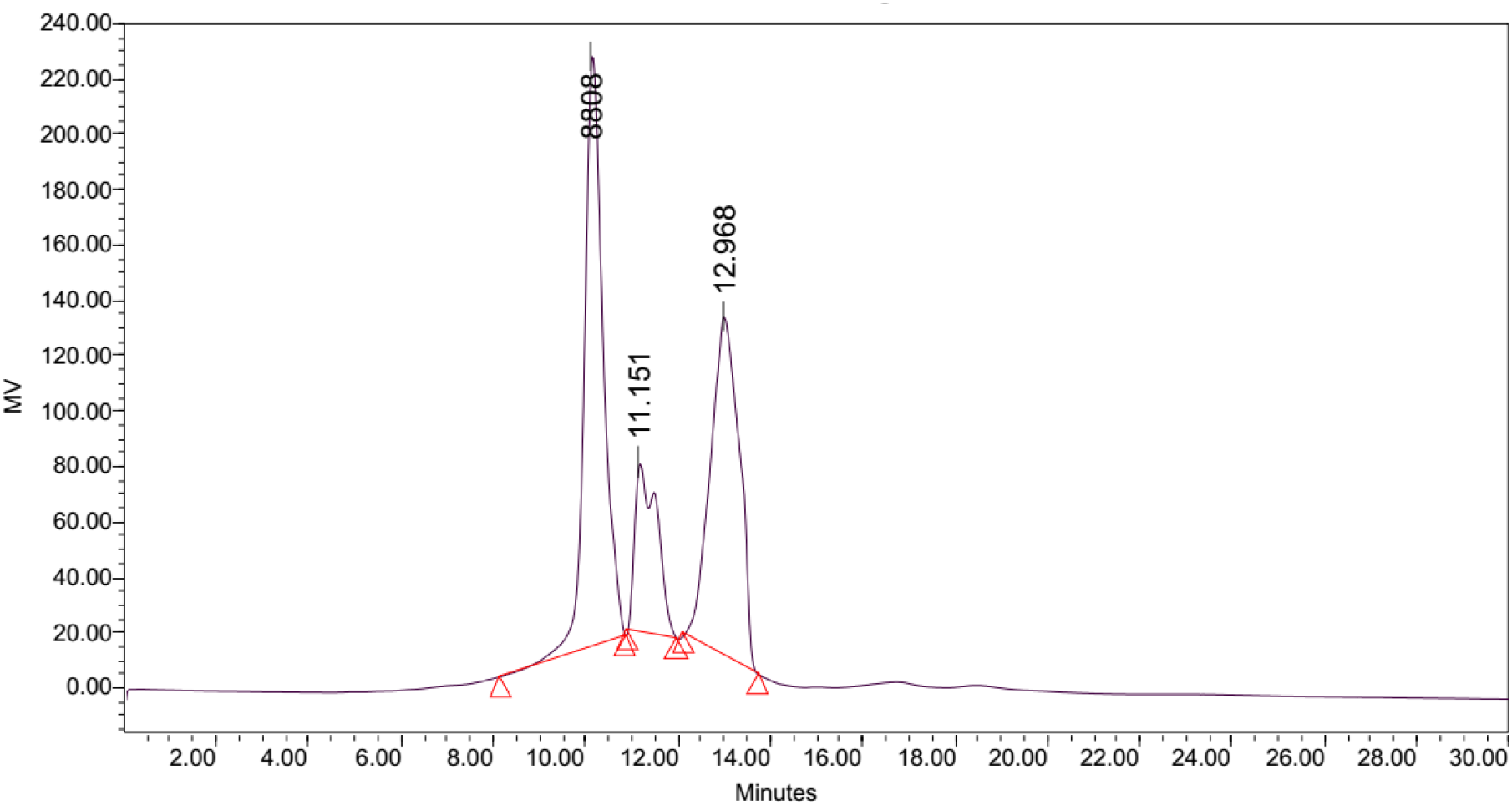
Gel filtration chromatography separation of LB Lennox broth at time = 1 hrs reveals the existence of three peaks. Injection volume = 10 μL, flow rate = 1 ml/min. Column: Agilent PL aquagel-OH MIX 8 μm, refractive index detector

**Figure 4:**
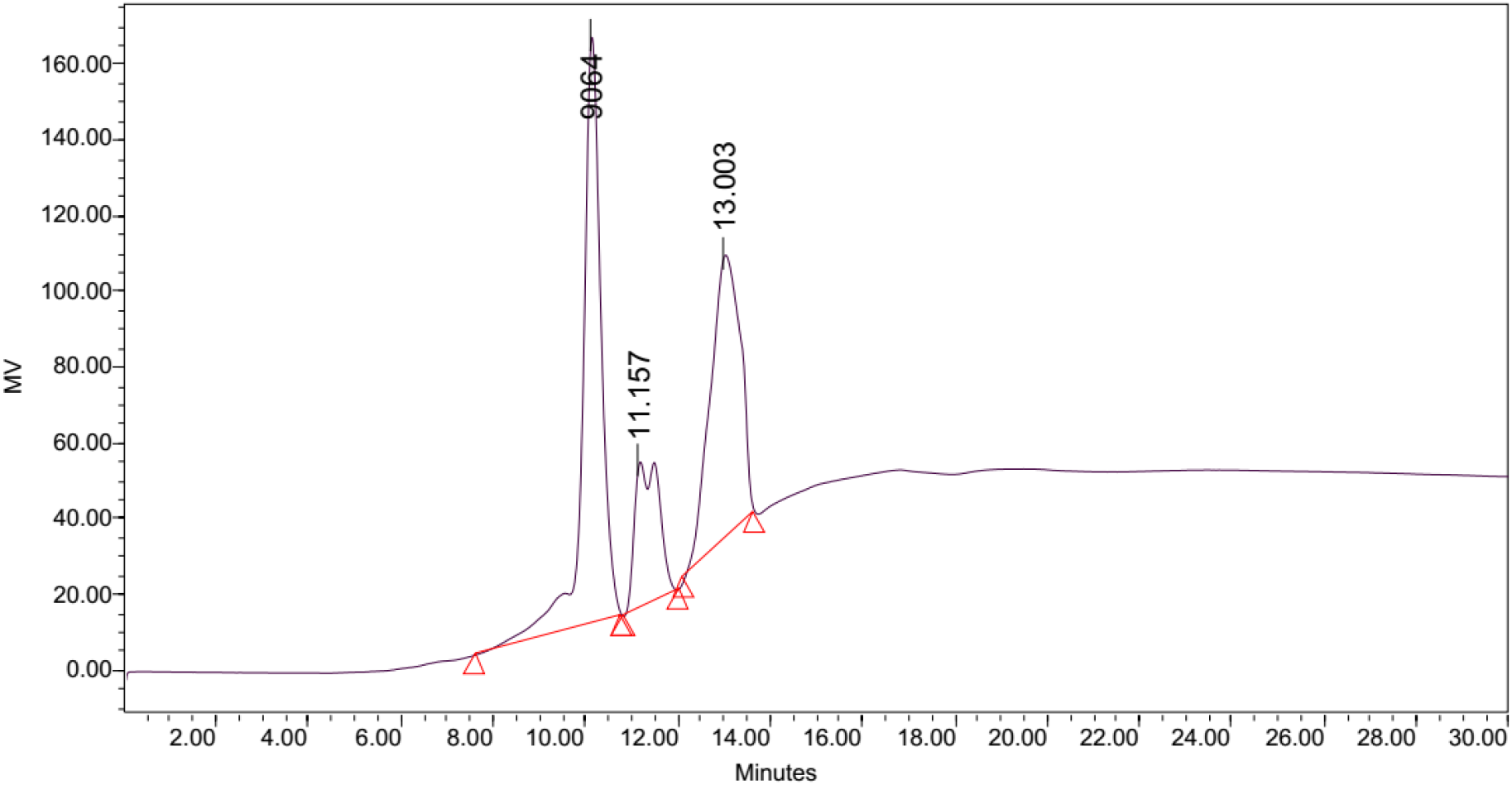
Gel filtration chromatography separation of LB Lennox broth at time = 2 hrs reveals the existence of three peaks. Injection volume = 10 μL, flow rate = 1 ml/min. Column: Agilent PL aquagel-OH MIX 8 μm, refractive index detector

**Figure 5:**
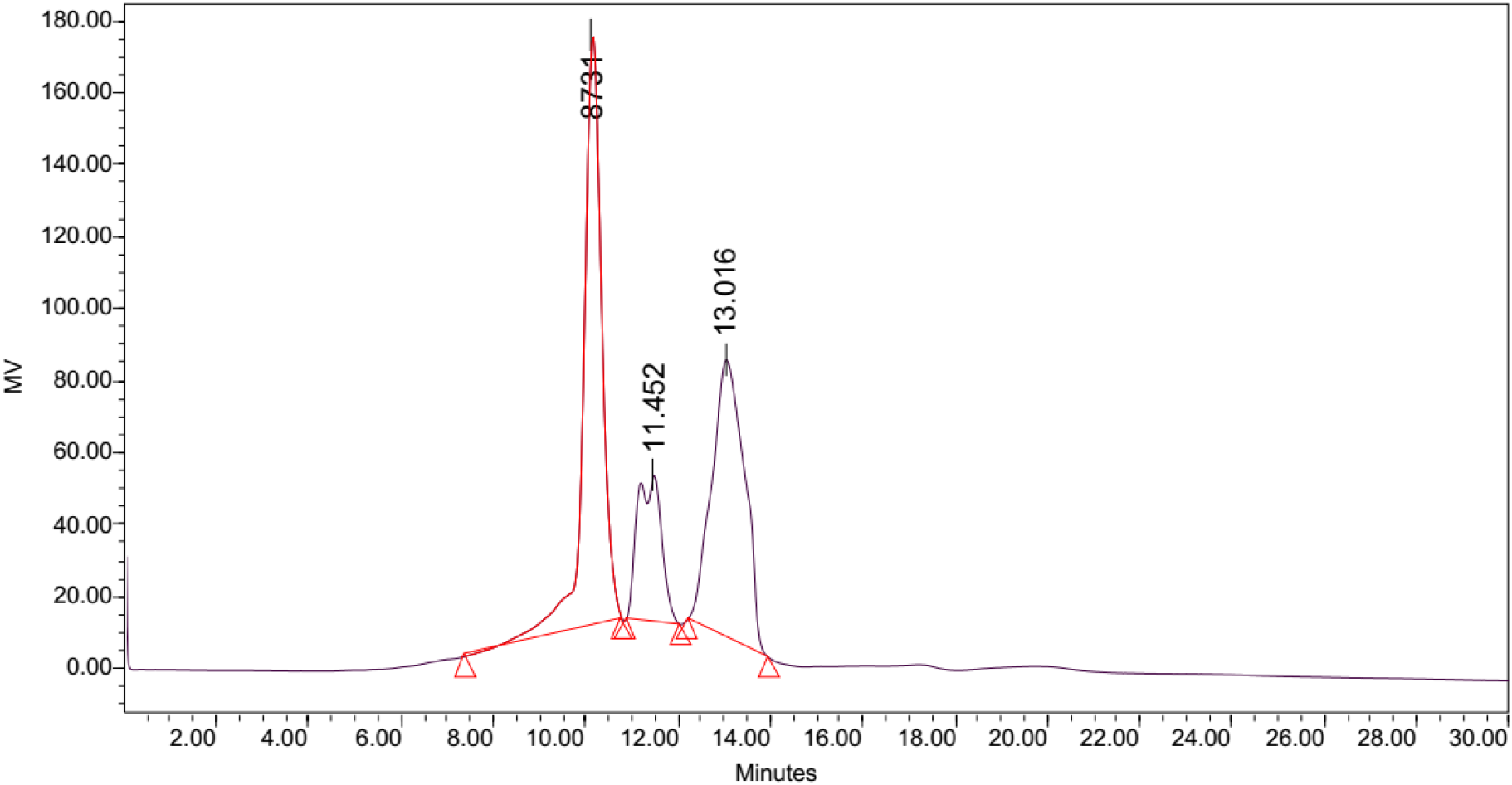
Gel filtration chromatography separation of LB Lennox broth at time = 3 hrs reveals the existence of three peaks. Injection volume = 10 μL, flow rate = 1 ml/min. Column: Agilent PL aquagel-OH MIX 8 μm, refractive index detector

**Figure 6:**
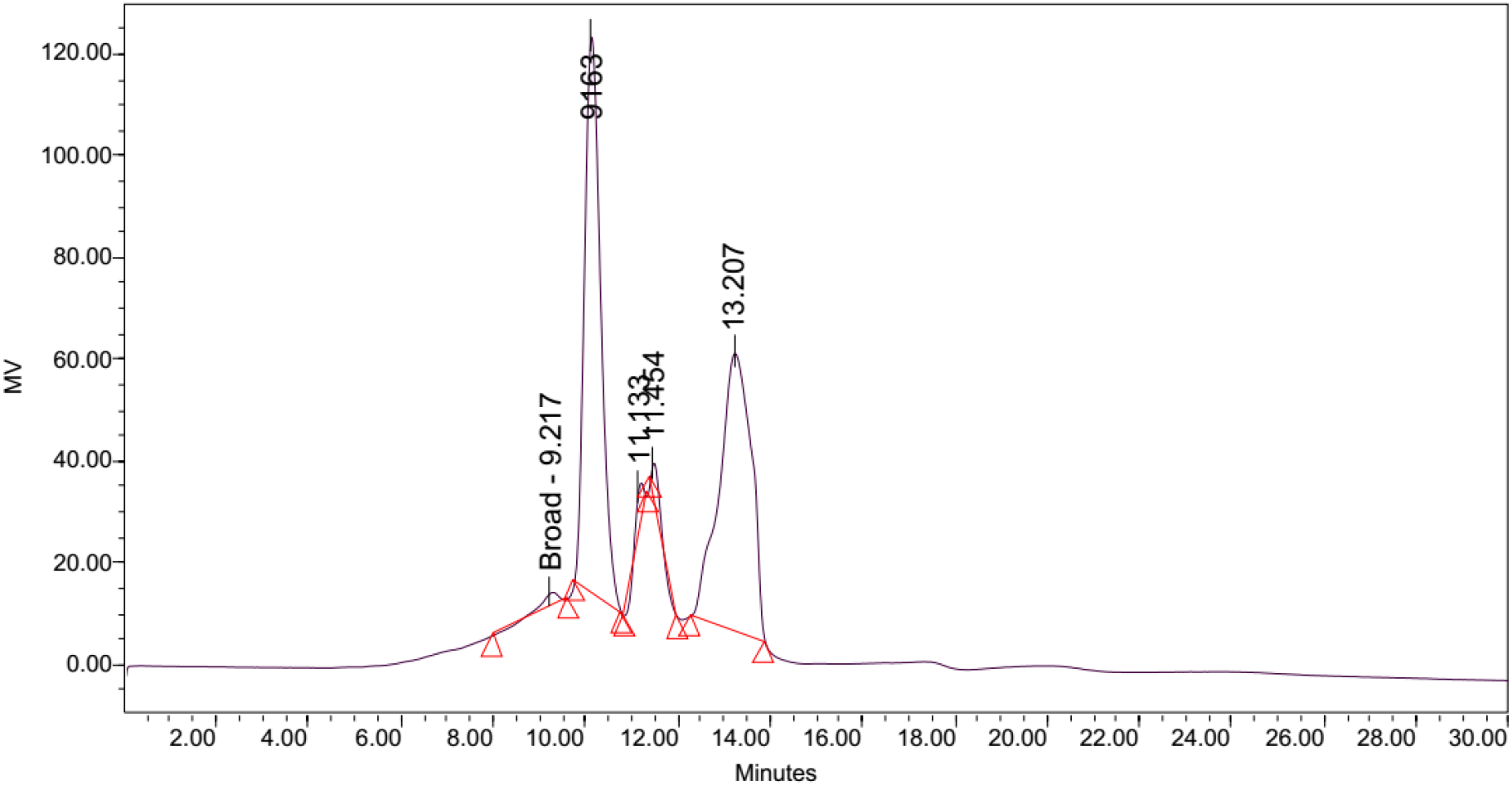
Gel filtration chromatography separation of LB Lennox broth at time = 4 hrs reveals the existence of four peaks. Injection volume = 10 μL, flow rate = 1 ml/min. Column: Agilent PL aquagel-OH MIX 8 μm, refractive index detector

**Figure 7:**
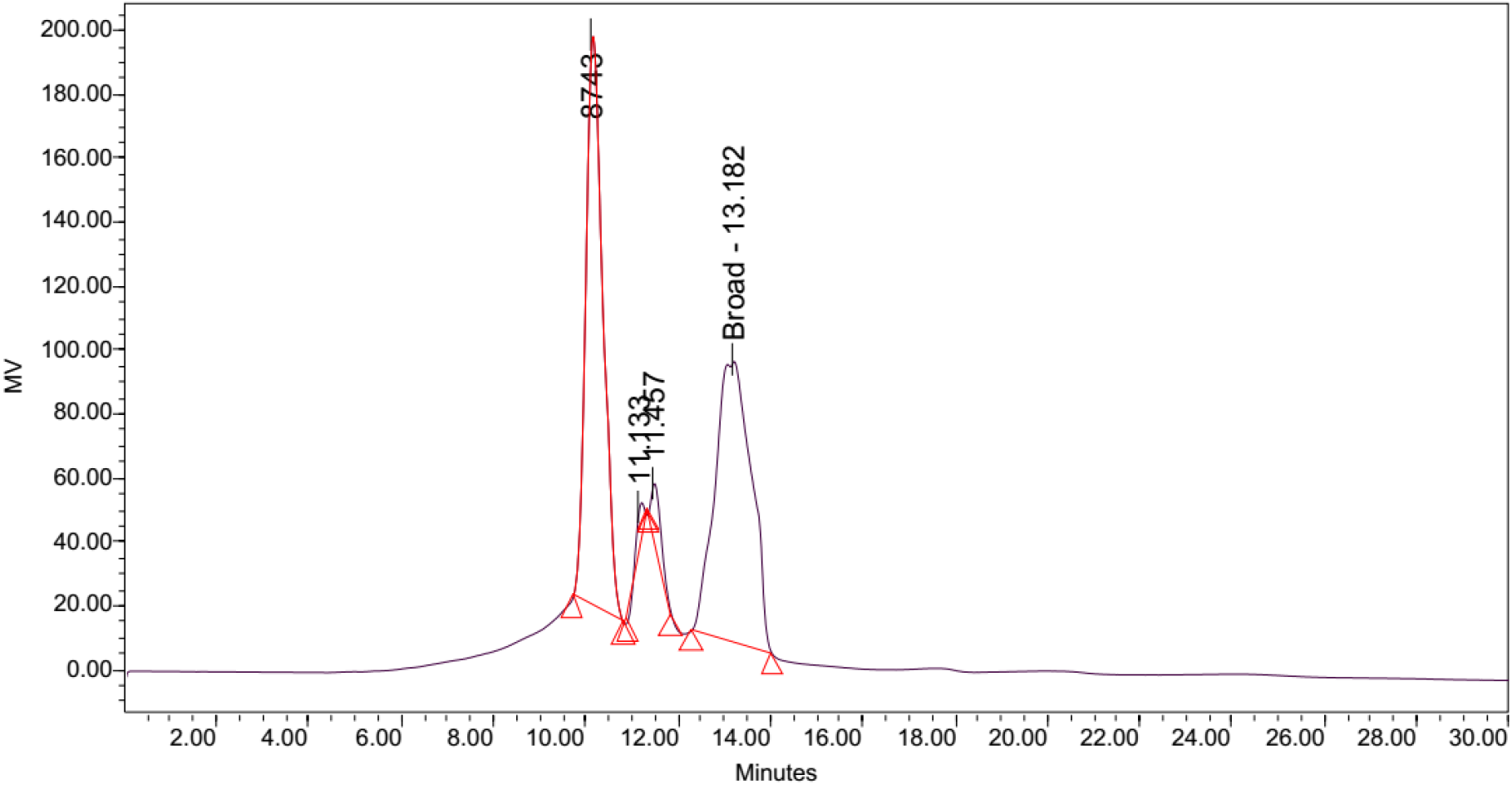
Gel filtration chromatography separation of LB Lennox broth at time = 5 hrs reveals the existence of three peaks. Injection volume = 10 μL, flow rate = 1 ml/min. Column: Agilent PL aquagel-OH MIX 8 μm, refractive index detector

**Figure 8:**
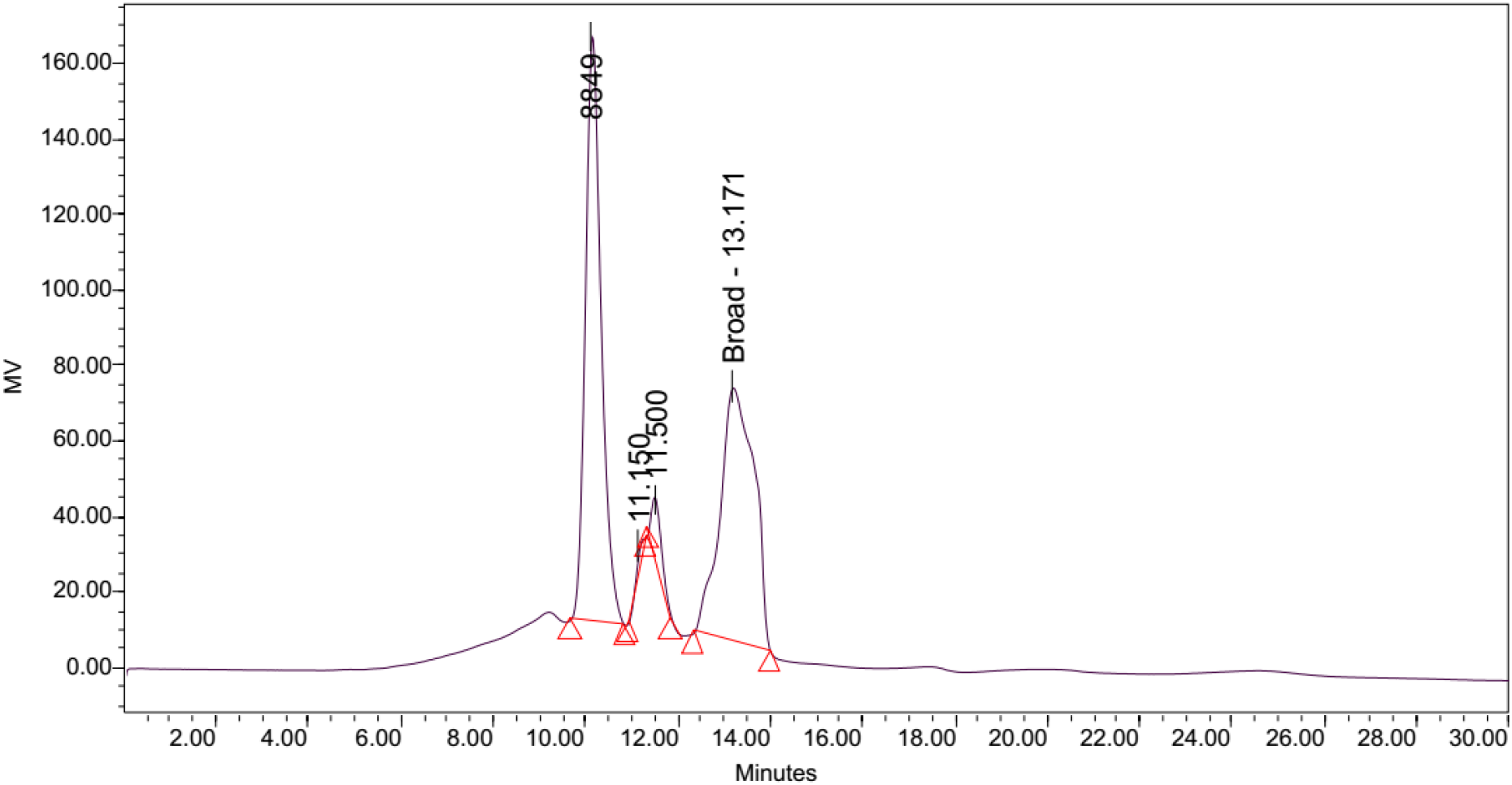
Gel filtration chromatography separation of LB Lennox broth at time = 6 hrs reveals the existence of three peaks. Injection volume = 10 μL, flow rate = 1 ml/min. Column: Agilent PL aquagel-OH MIX 8 μm, refractive index detector

Given that wavelength for visualizing a chromatogram plays a critical role in determining the relative success of different elution conditions and gradient profile, 194 nm visualization of reversed phase high performance liquid chromatography (RP-HPLC) was selected in preference to 280 nm (information from Figure S6, S7 and S8).

Elution of LB Lennox broth at 1 hour post inoculation from C-18 RP-HPLC is shown in Figure 9, which indicated the complexity of the broth that precludes clean separation by RP-HPLC, as well as the presence of large amounts of hydrophilic compounds which could be desorbed by a 5% ethanol/95% water mixture. Similarly, Figure 10 reveals a broth mixture at 6 hours post inoculation that could not be separated into different fractions by RP-HPLC. Comparisons between the different visualization wavelengths of 280, 264, 254, 194 and 380 nm in diode array detection of chromatogram reveals that fewer peaks were present in chromatograms visualized at 280, 264, and 254 nm compared to 194 nm (Figure S6, S7 and S8), which validated the use of 194 nm as the visualization wavelength for the current fractionation experiment involving LB Lennox broth. Visualization at 380 nm did not yield appreciable peaks (Figure S6, S7 and S8).

**Figure 9:**
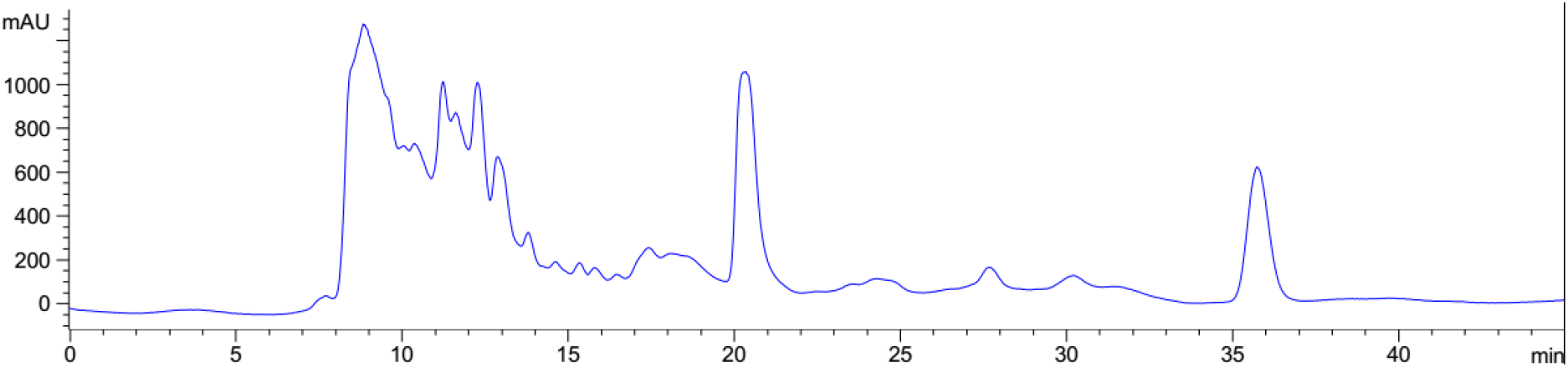
Elution profile of LB Lennox broth at 1 hour post inoculation with RP-HPLC. Detection by diode array detector at 194 nm, Column: Agilent Technologies Poroshell 120 SB-C18, Flow rate = 0.2 ml/min, 5/95 Ethanol/water (%/%) for isocratic elution, 25 °C, 70 bar pressure, Injection volume = 10 μL

**Figure 10:**
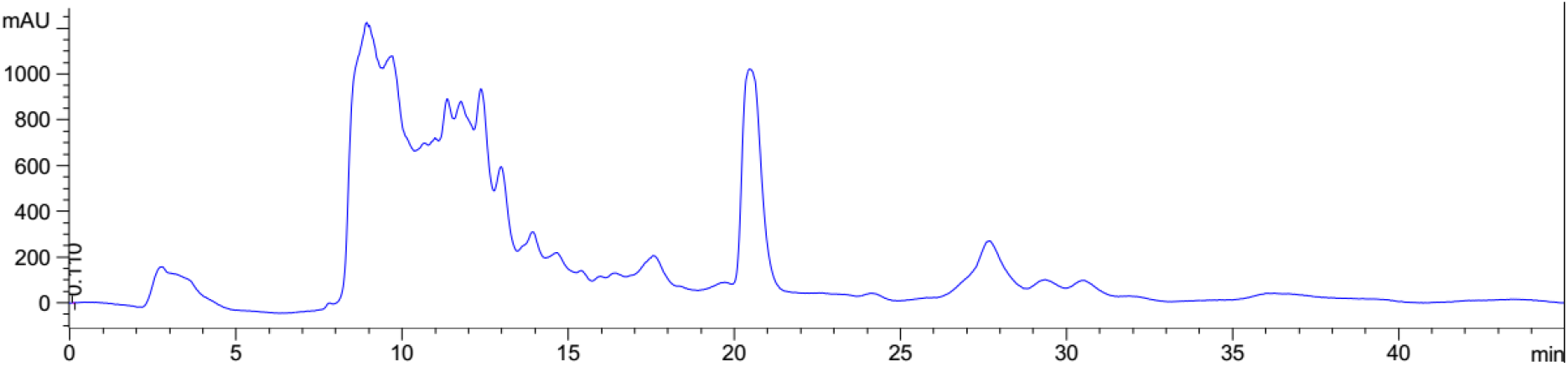
Elution profile of LB Lennox broth at 6 hour post inoculation with RP-HPLC. Detection by diode array detector at 194 nm, Column: Agilent Technologies Poroshell 120 SB-C18, Flow rate = 0.2 ml/min, 5/95 Ethanol/water (%/%) for isocratic elution, 25 °C, 70 bar pressure, Injection volume = 10 μL

Application of a gradient elution profile did not significantly improve the fractionation of LB Lennox broth at 1 hour post inoculation (Figure 11). Specifically, the eluted chromatogram revealed no fractionation of the complex mixture into a couple of distinct fractions.

**Figure 11:**
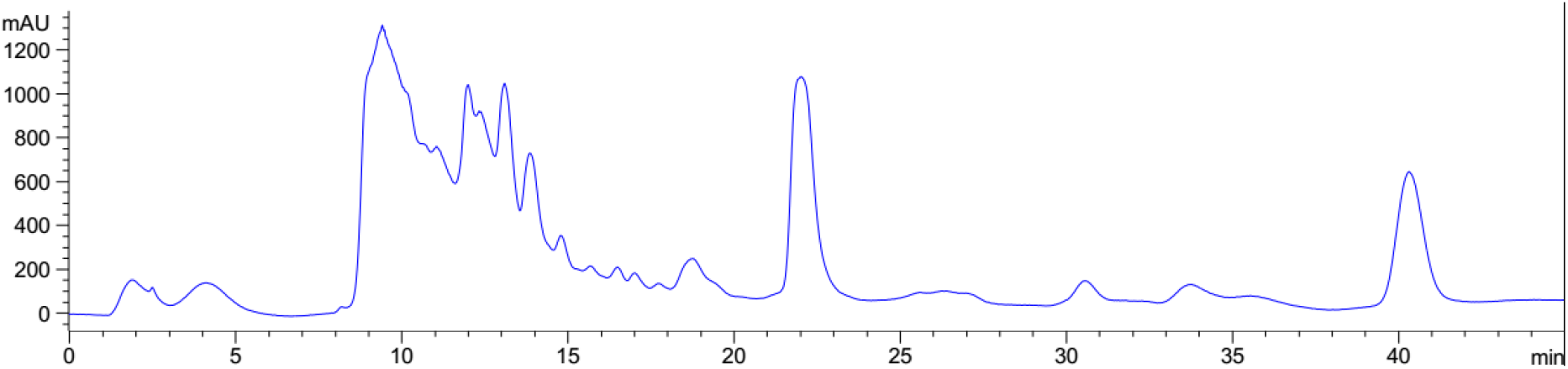
Elution profile of LB Lennox broth at 1 hour post inoculation with RP-HPLC. Detection by diode array detector at 194 nm, Column: Agilent Technologies Poroshell 120 SB-C18, Flow rate = 0.2 ml/min, 5/95 Ethanol/water (%/%) for gradient elution, 25 °C, 68 bar pressure, Injection volume = 10 μL

Use of a linear gradient with 1% increase in ethanol/water ratio per minute did not result in better fractionation of the LB Lennox broth mixture at 1 hr post inoculation as visualized on 194 nm wavelength (Figure 12). Specifically, separation worsened due probably to the lack of equilibration between the column and the mobile phase.

**Figure 12:**
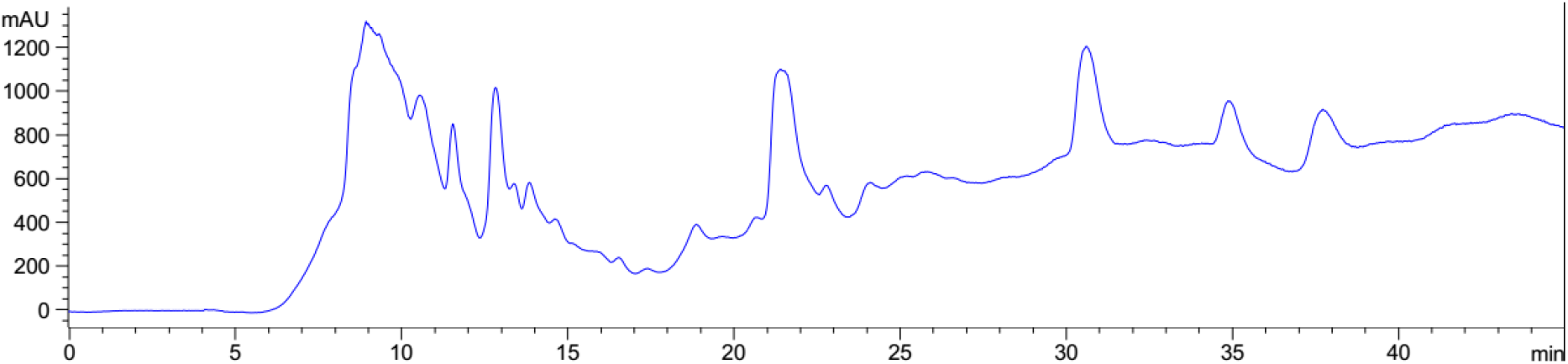
Elution profile of LB Lennox broth at 1 hour post inoculation with RP-HPLC. Detection by diode array detector at 194 nm, Column: Agilent Technologies Poroshell 120 SB-C18, Flow rate = 0.2 ml/min, linear gradient with 1% increase in ethanol/water ratio per min, t=0, 5% ethanol, maximum = 40% ethanol, 25 °C, 70 bar pressure, Injection volume = 10 μL

## 4. Conclusions

Determining the compositional changes in growth medium that accompany cell growth is a grand challenge in microbial physiology studies, especially in the case where complex chemically undefined medium is used. Given the multitude of compounds and components present, separation of the growth medium mixture into distinct components remain impossible, despite the advent of refinements in high performance liquid chromatography methods. Motivated by the desire to visualize the likely changes in molecular weight and components which growth of *Escherichia coli* DH5α (ATCC 53868) brought forth to LB Lennox medium, this study used a combination of gel filtration chromatography (GFC) and C-18 reversed phase high performance liquid chromatography (RP-HPLC) for fractionating the complex broth mixture. Results from GFC characterization of the molecular weight changes during cultivation of *E. coli* DH5α did not capture any salient details due likely to the inability of the column used in resolving molecular weight changes on the order of a few hundreds daltons. On the other hand, attempted fractionation of LB Lennox broth through various elution profiles on RP-HPLC failed to reveal distinct fractions at 194 nm detection wavelength, which, on the experience of this study, should be used for visualizing the chromatogram resulting from separation of LB Lennox medium. Collectively, although the study failed to highlight molecular weight changes in LB Lennox broth at the component level, it nevertheless validated the use of 194 nm detection wavelength as the preferred visualization wavelength for understanding the elution profile of LB Lennox from a C-18 RP-HPLC analytical run. Additionally, results obtained indicated the significant challenge involved in separating or fractionating a microbiological medium.

## Supporting information

Supplementary materials of manuscript

## Supplementary materials

The supplementary file appended with this manuscript contains chromatograms of calibration of gel filtration chromatography as well as reversed phase high performance liquid chromatography analysis of LB Lennox broth at different timepoints in the bioreactor cultivation of *Escherichia coli* DH5α.

## Author’s contribution

The author developed the idea and concept for the experiment, performed the experiment, analysed the data, and wrote the manuscript.

