## Supplementary materials of manuscript for "Understanding the compositional changes in LB Lennox medium during growth of *Escherichia coli* DH5α in a 1L bioreactor"

Wenfa Ng

Department of Chemical and Biomolecular Engineering, National University of Singapore

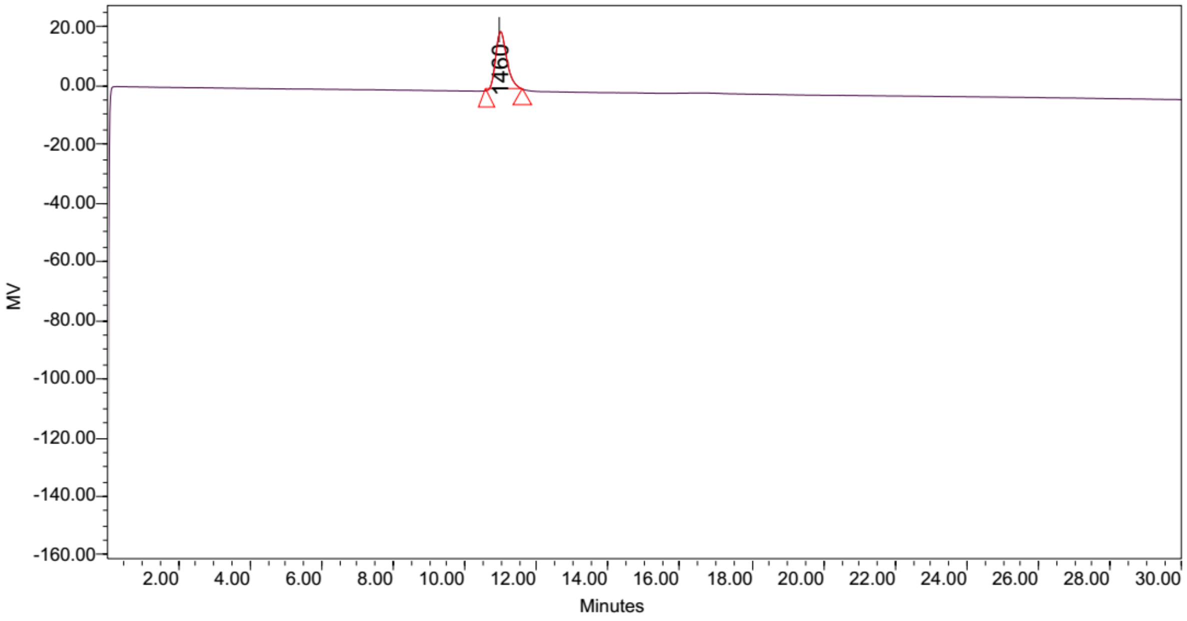


**Figure S1:** Elution of calibration standard 1, molecular weight = 1460 Da from Waters Gel filtration chromatography instrument with Agilent PL aquagel-OH MIX 8 µm column. Flow rate = 1 ml/min, Injection volume = 10 µL, Refractive index detector

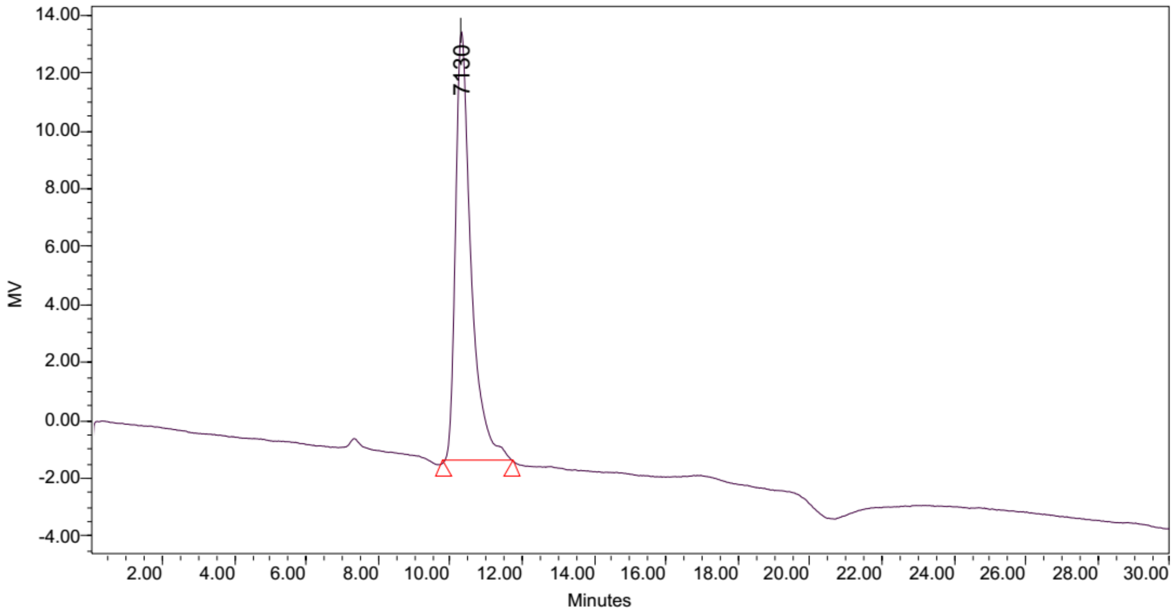


**Figure S2:** Elution of calibration standard 2, molecular weight = 7130 Da from Waters Gel filtration chromatography instrument with Agilent PL aquagel-OH MIX 8 µm column. Flow rate = 1 ml/min, Injection volume = 10 µL, Refractive index detector

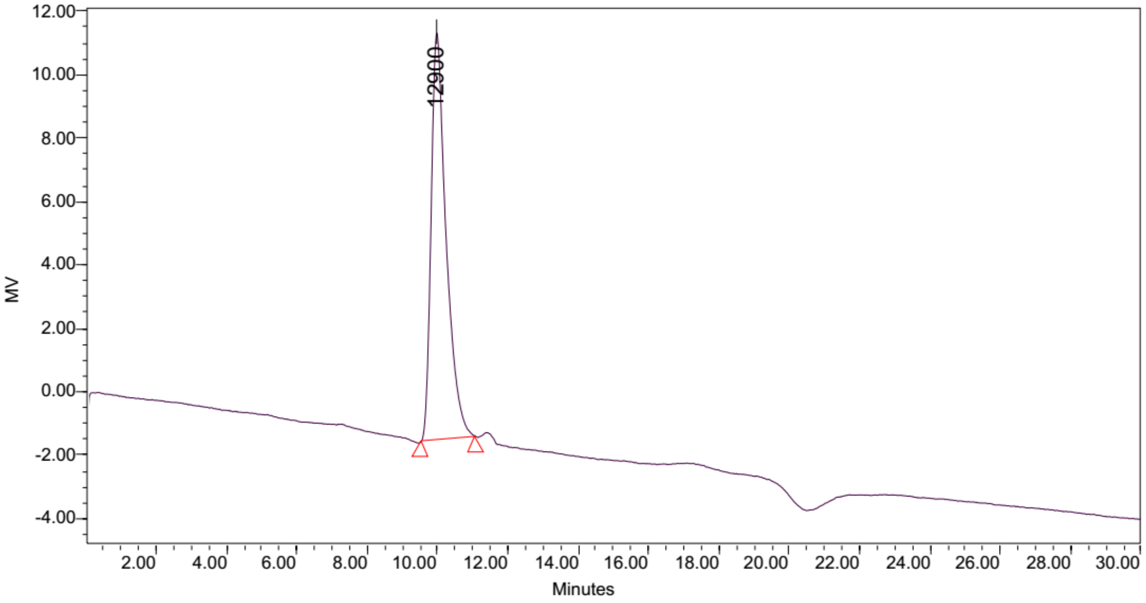


**Figure S3:** Elution of calibration standard 3, molecular weight = 12900 Da from Waters Gel filtration chromatography instrument with Agilent PL aquagel-OH MIX 8 µm column. Flow rate = 1 ml/min, Injection volume = 10 µL, Refractive index detector

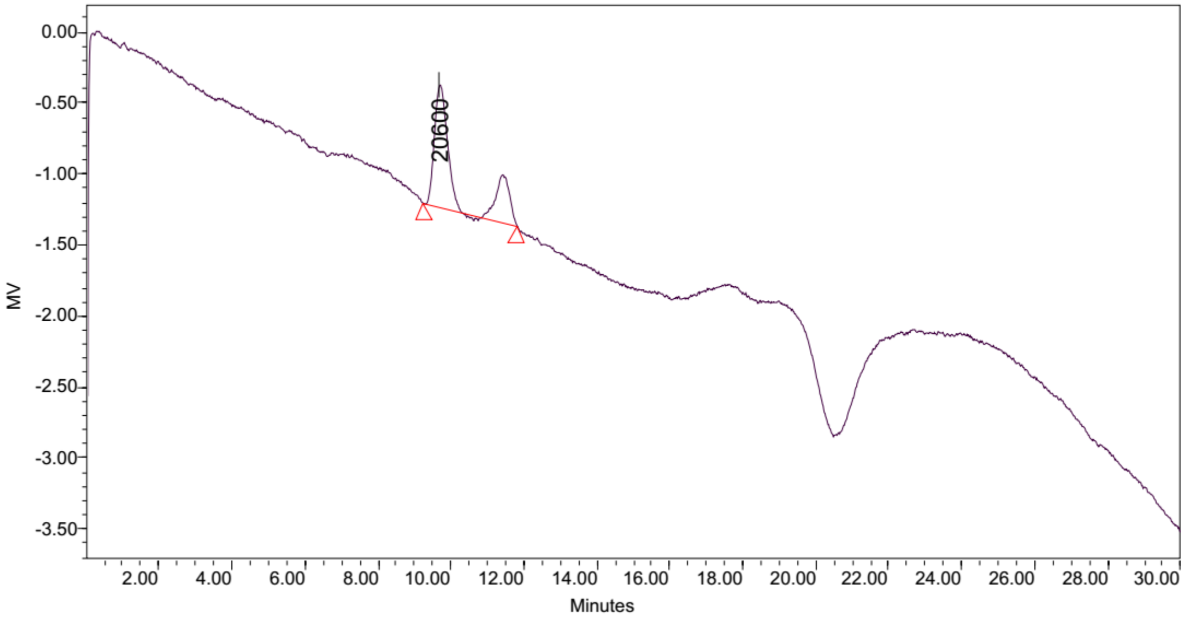


**Figure S4:** Elution of calibration standard 4, molecular weight = 20600 Da from Waters Gel filtration chromatography instrument with Agilent PL aquagel-OH MIX 8 µm column. Flow rate = 1 ml/min, Injection volume = 10 µL, Refractive index detector

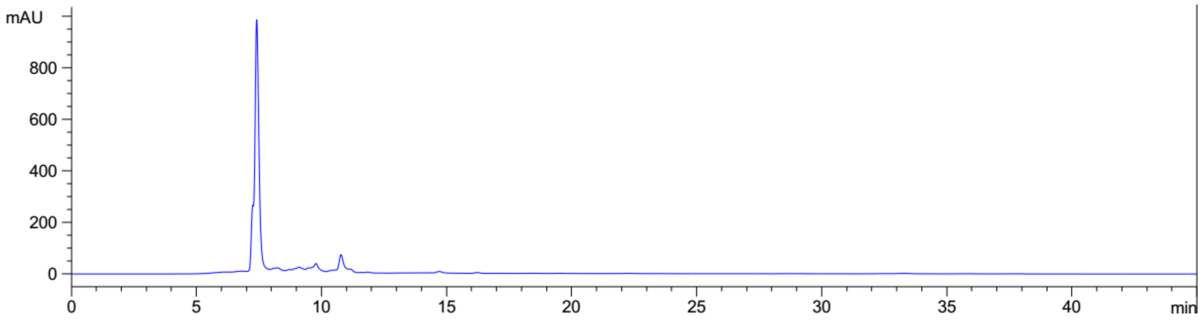


**Figure S5:** Elution profile of LB Lennox broth at 6 hours post inoculation with RP-HPLC. Detection by variable wavelength detector at 280 nm, Column: Agilent Technologies Poroshell 120 SB-C18, Flow rate = 0.2 ml/min, 5/95 Ethanol/water (%/%) for isocratic elution, 25 $^{o}$C, 81 bar pressure, Injection volume = 10 µL

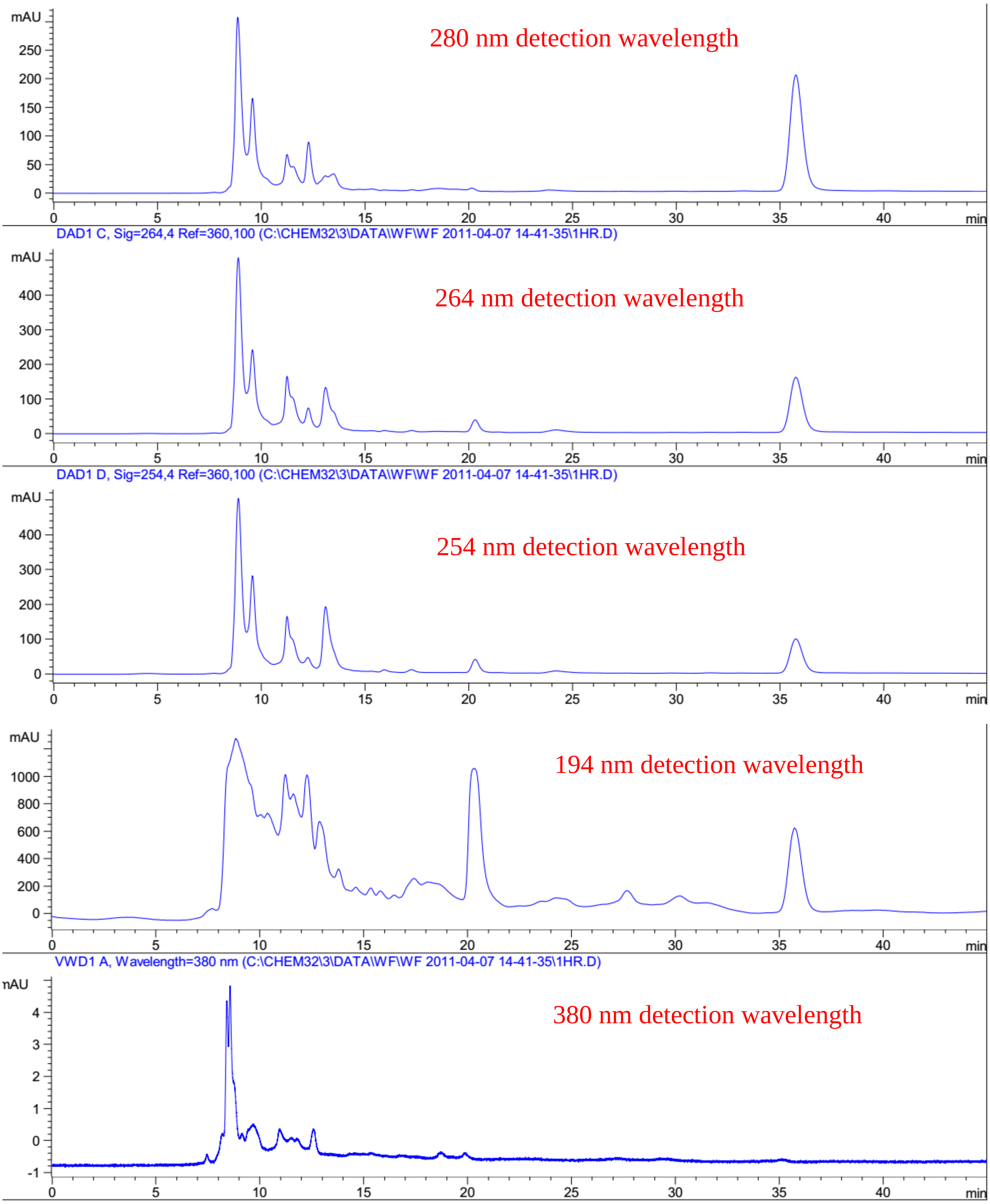


**Figure S6:** Elution profile of LB Lennox broth at 1 hour post inoculation with RP-HPLC. Detection by diode array detector, Column: Agilent Technologies Poroshell 120 SB-C18, Flow rate = 0.2 ml/min, 5/95 Ethanol/water (%/%) for isocratic elution, 25 $^{o}$C, 70 bar pressure, Injection volume = 10 µL

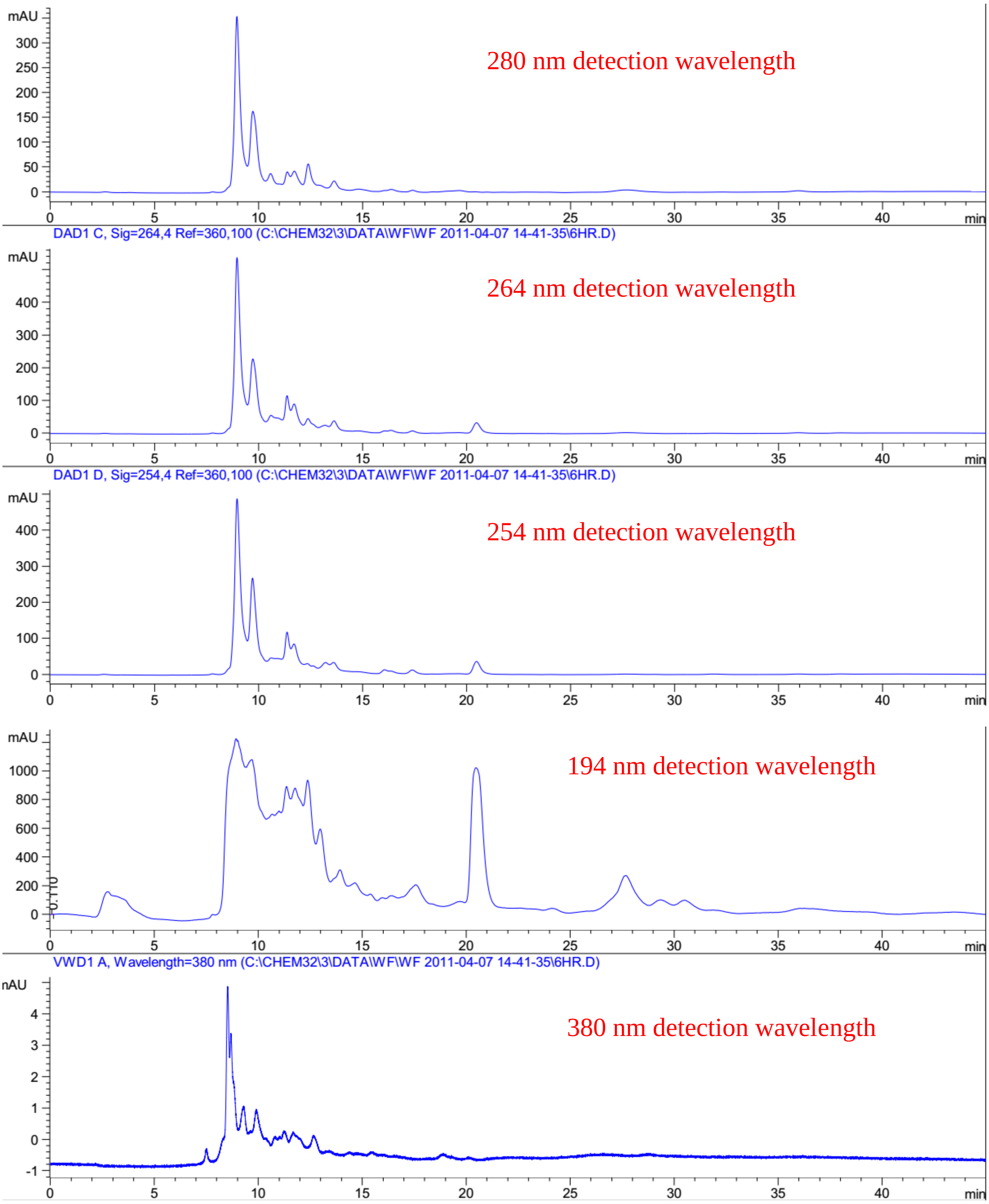


**Figure S7:** Elution profile of LB Lennox broth at 6 hour post inoculation with RP-HPLC. Detection by diode array detector, Column: Agilent Technologies Poroshell 120 SB-C18, Flow rate = 0.2 ml/min, 5/95 Ethanol/water (%/%) for isocratic elution, 25 $^{o}$C, 70 bar pressure, Injection volume = 10 µL

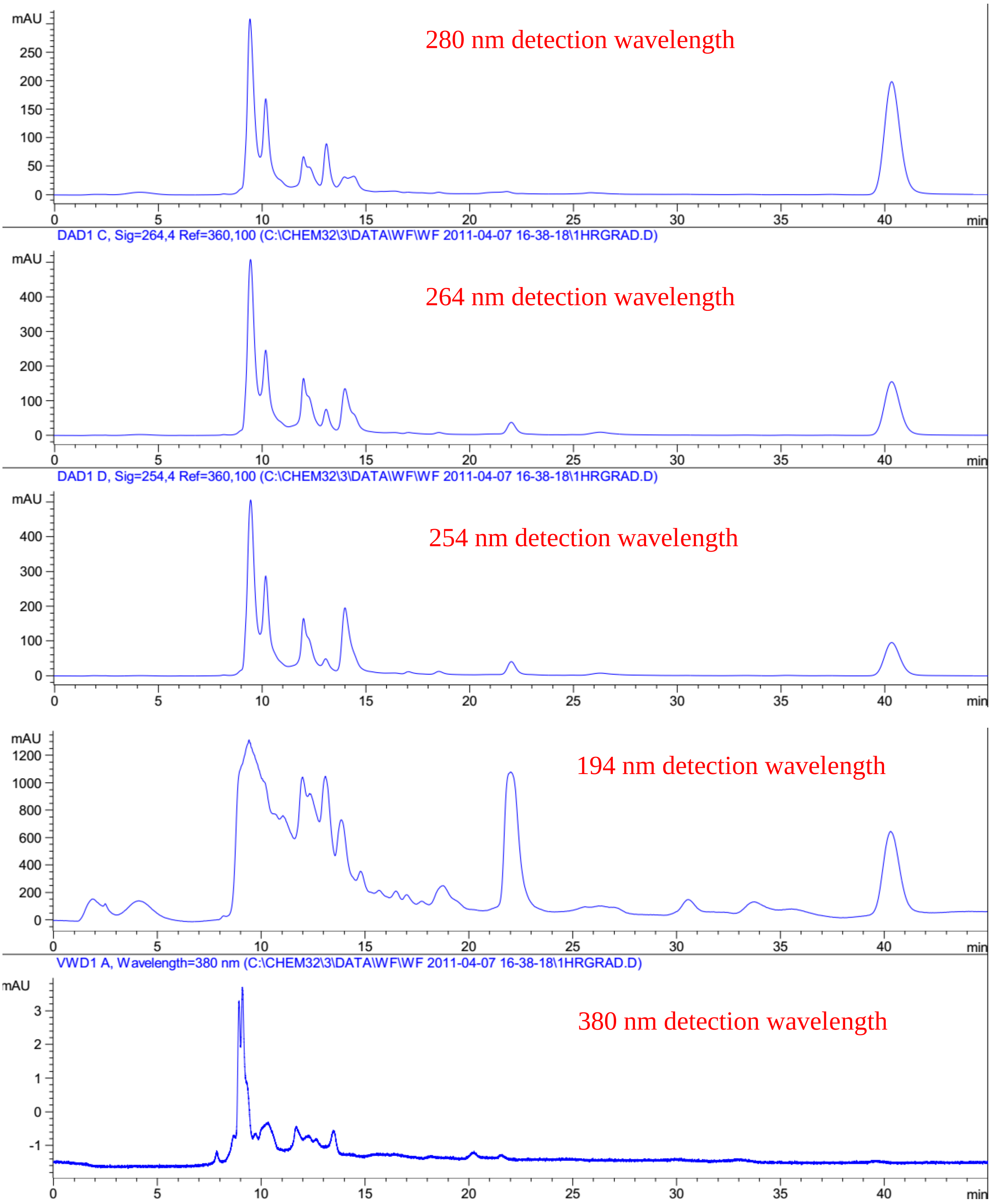


**Figure S8:** Elution profile of LB Lennox broth at 1 hour post inoculation with RP-HPLC. Detection by diode array detector, Column: Agilent Technologies Poroshell 120 SB-C18, Flow rate = 0.2 ml/min, 5/95 Ethanol/water (%/%) for gradient elution, 25 $^{o}$C, 68 bar pressure, Injection volume = 10 µL

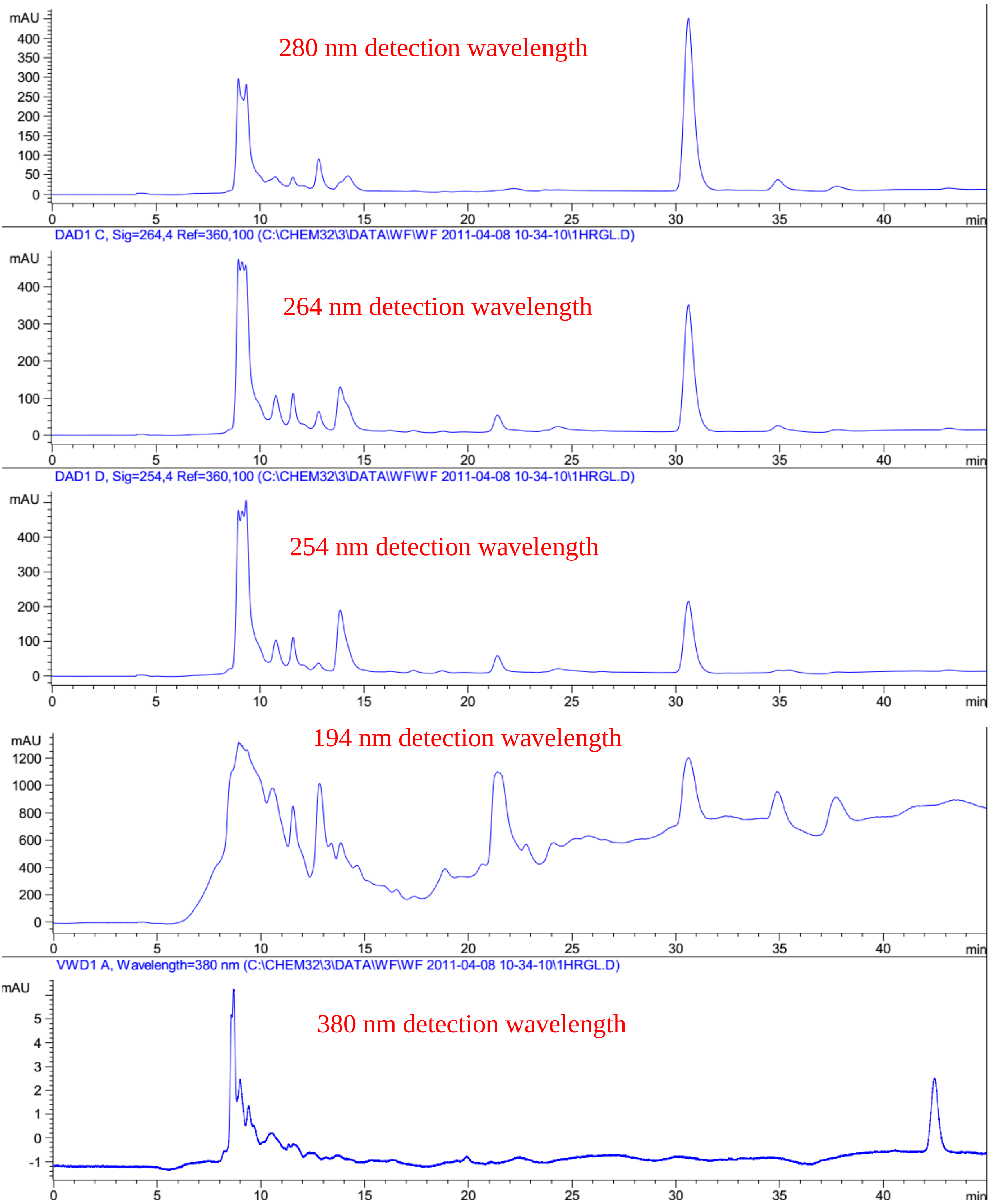


**Figure S9:** Elution profile of LB Lennox broth at 1 hour post inoculation with RP-HPLC. Detection by diode array detector, Column: Agilent Technologies Poroshell 120 SB-C18, Flow rate = 0.2 ml/min, linear gradient with 1% increase in ethanol/water ratio per min, t=0, 5% ethanol, maximum = 40% ethanol, 25 °C, 70 bar pressure, Injection volume = 10 μL
